## Supplementary figures and table for "Allosteric coupling asymmetry mediates paradoxical activation of BRAF by type II inhibitors"

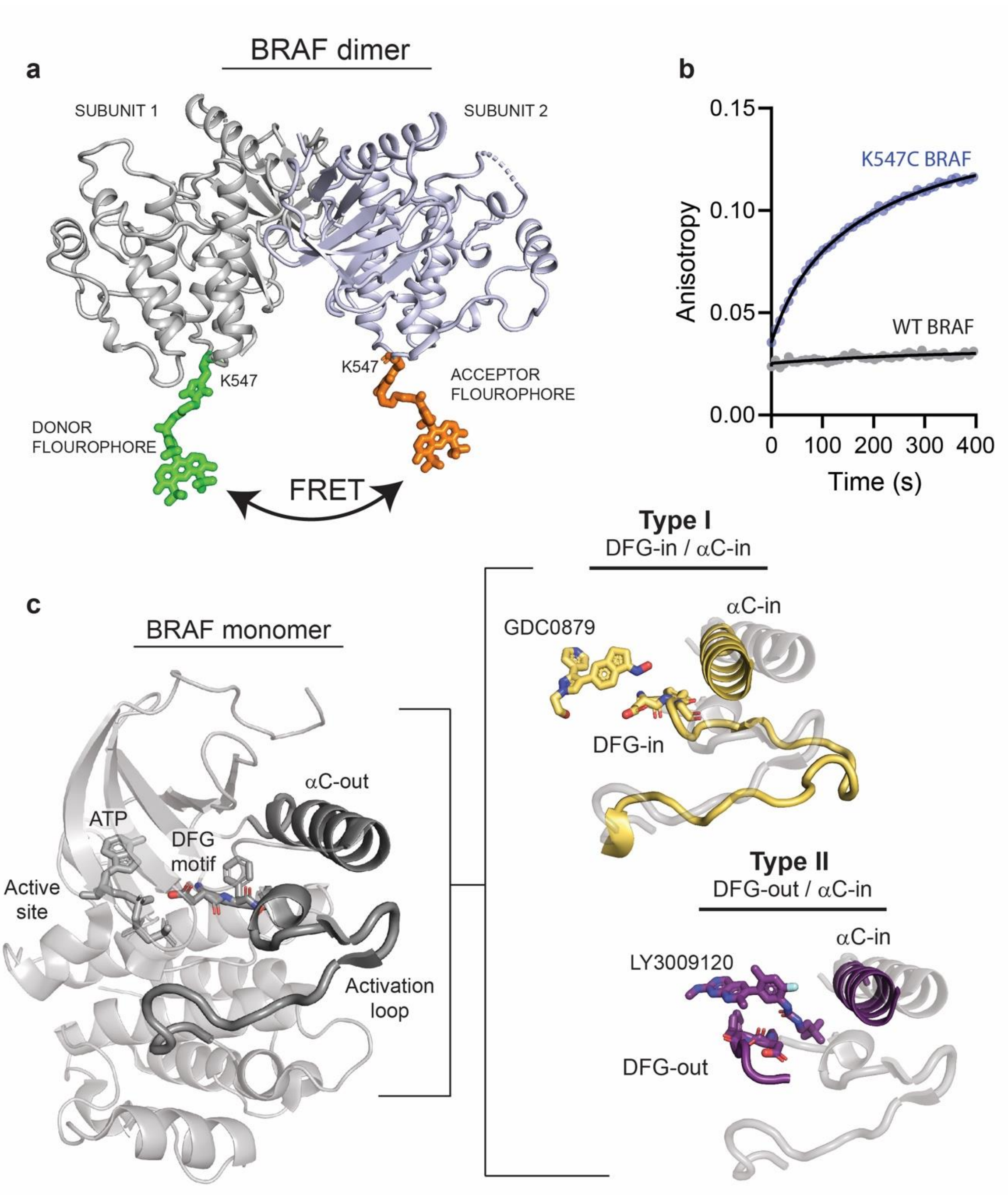


**Supplementary Figure 1. X-ray structures showing the BRAF dimer labeled with fluorophores and structural features of type I and type II inhibitor binding modes. a)** X-ray structure (PDB: 5C9C) showing the BRAF dimer covalently labeled at K547 with Alexa Fluor 488 C5-maleimide using mtsslSuite (http://www.mtsslsuite.isb.ukbonn.de/). **b**) Stopped flow fluorescence anisotropy data showing labeling of BRAF WT and K547C with Alexa 488. **c)** X-ray structure of the BRAF monomer (PDB ID: 6PP9) highlighting (dark grey) key regulatory features of the kinase domain including the DFG-motif, αC-helix, and activation loop. The structural differences that distinguish type I (yellow) and type II (purple) inhibitors are highlighted and overlaid with the structure of the autoinhibited BRAF monomer (light grey) (PDB IDs: 4MNE, 5C9C).

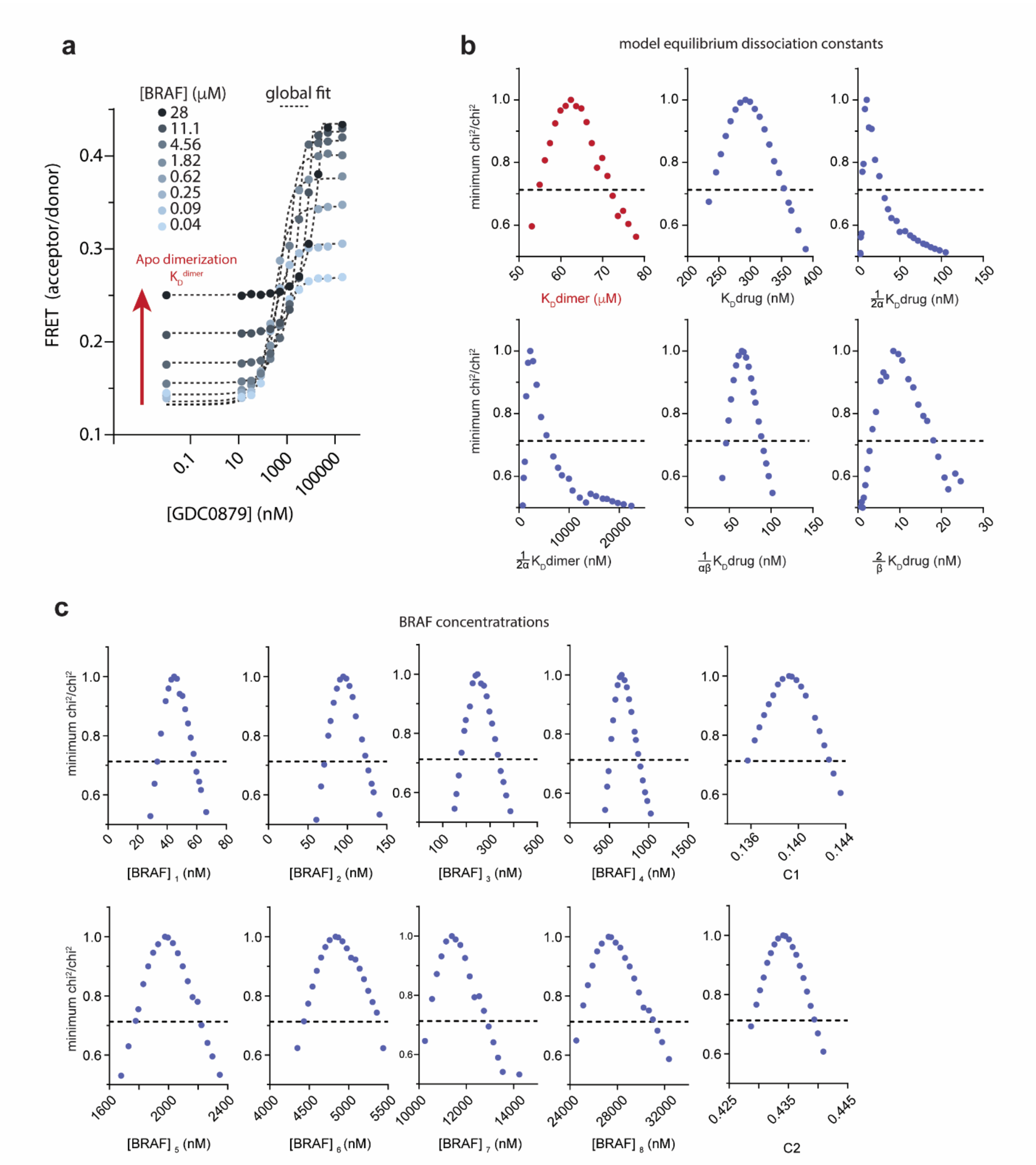


**Supplementary Figure 2. Quantifying apo BRAF dimerization affinity ($K_D^{dimer}$). a)** Intermolecular FRET experiments tracking BRAF dimerization as a function of GDC0879 concentration were carried out at high BRAF concentrations to define the relatively weak apo BRAF dimerization affinity. Black dotted lines represent a global fit to the thermodynamic model shown in Figure 1c (see Methods). The red arrow highlights apo BRAF dimerization ($K_D^{dimer}$) occurring with no inhibitor. **b,c)** Representative one-dimensional error surface analysis of the global fit parameters for the FRET data shown in panel a, including equilibrium dissociation constants (**b**) and BRAF concentrations and fluorescence coefficients (**c**). The black dotted line represents the chi$^2$ threshold used to establish 95% CIs for each parameter (see Methods). Note that all parameters shown are well-constrained within this limit as indicated by their intersection with the chi$^2$ threshold boundary.

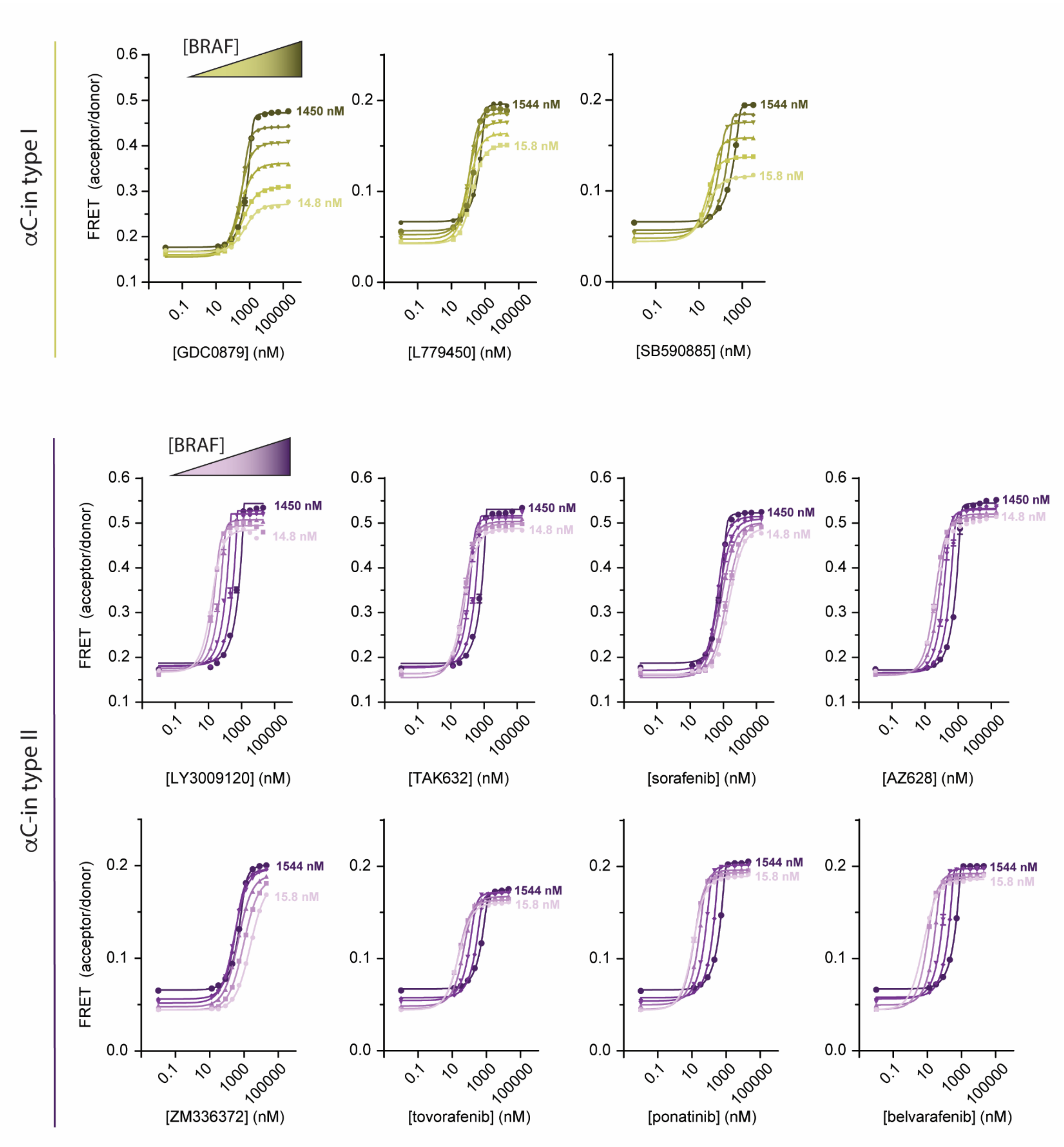


**Supplementary Figure 3. Intermolecular FRET experiments measuring inhibitor-induced BRAF dimerization.** Representative data for all αC-in inhibitors showing BRAF dimerization for type I (yellow) and type II (purple) inhibitors. Gradients from light to dark represent experiments done at increasing BRAF concentrations. Lines are fits of individual titrations to a quadratic binding model in Graphpad prism and are for illustrative purposes only; all data were globally fit to the thermodynamic model shown in Figure 1c.

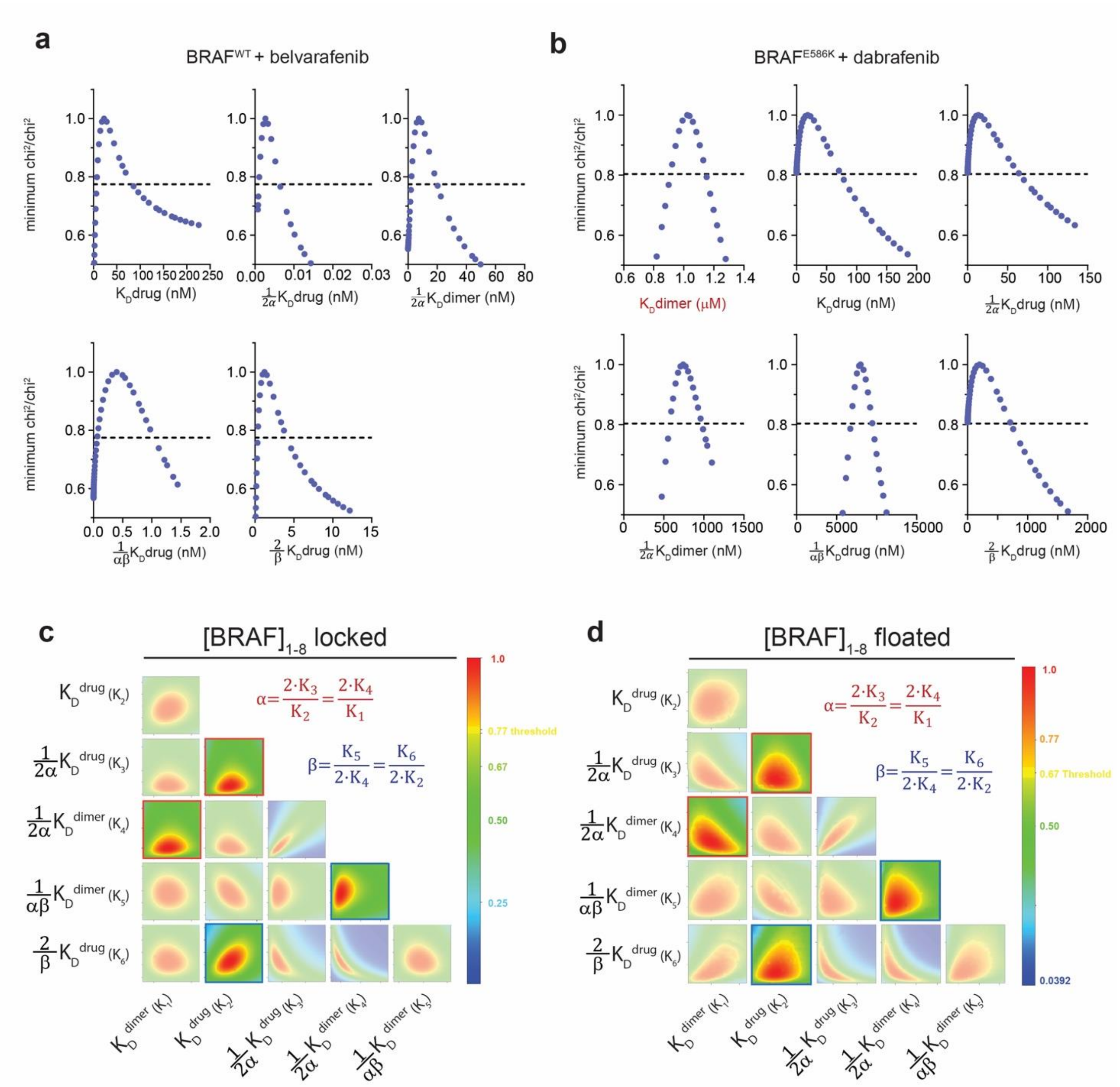


**Supplementary Figure 4. One- and two-dimensional error surfaces show that global fitting yields well-constrained parameters.** Representative one-dimensional error surface analyses of the global fit parameters from FRET experiments with the $\alpha$C-in type II inhibitor belvarafenib (**a**) and the $\alpha$C-out inhibitor dabrafenib (**b**)**.** The black dotted line represents the chi$^2$ threshold used to establish 95% CIs for each parameter (see Methods). Note that all parameters shown are well-constrained within this limit as indicated by their intersection with the chi$^2$ threshold boundary. **c**,**d)** Representative two-dimensional confidence contour analyses of GDC0879 experiment shown in Supplementary Figure 2a, performed either with BRAF concentrations locked to their experimentally determined values (c) or allowed to float during fitting (d). All pair-wise combinations of unconstrained parameters were systematically varied to test for the presence of well-defined chi$^2$ minima and examine covariance between parameters. The color bar shows the chi$^2$ ratio with red representing the overall best-fit and yellow representing the chi$^2$ threshold boundary for 95% CIs. This analysis shows that all equilibrium constants as well as the allosteric coupling factors $\alpha$ and $\beta$, defined by the ratios of specific equilibrium dissociation constants (highlighted boxes, see Methods), are well-constrained in both fit procedures.

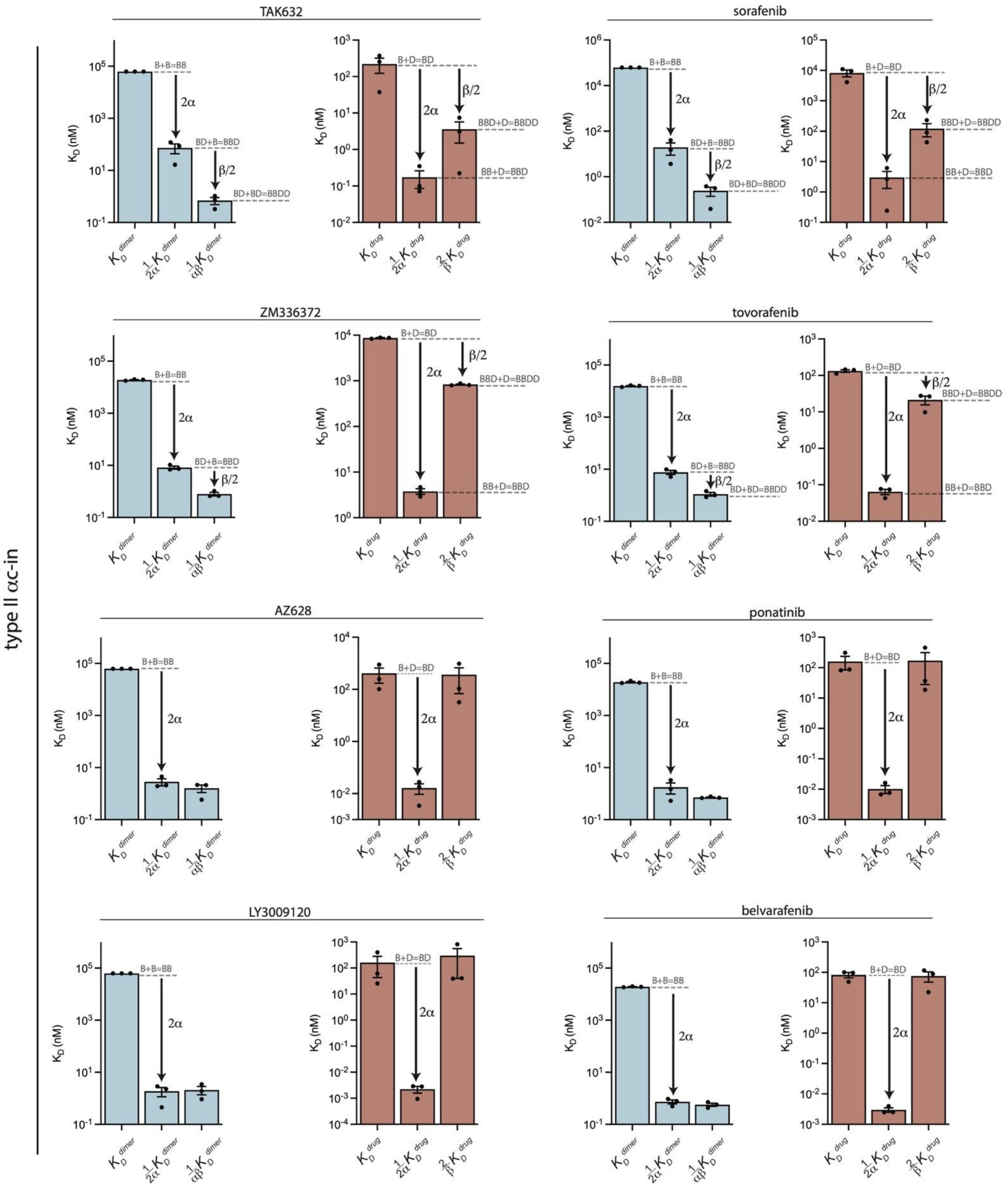
type II αc-in
TAK632
sorafenib
ZM336372
tovorafenib
AZ628
ponatinib
LY3009120
belvarafenib
K_D (nM)
B+B=BB
BD+B=BBD
BD+BD=BBDD
B+D=BD
BBD+D=BBDD
BB+D=BBD
2α
β/2
K_D^dimer
1/2α K_D^dimer
1/αβ K_D^dimer
K_D^drug
1/2α K_D^drug
2/β K_D^drug

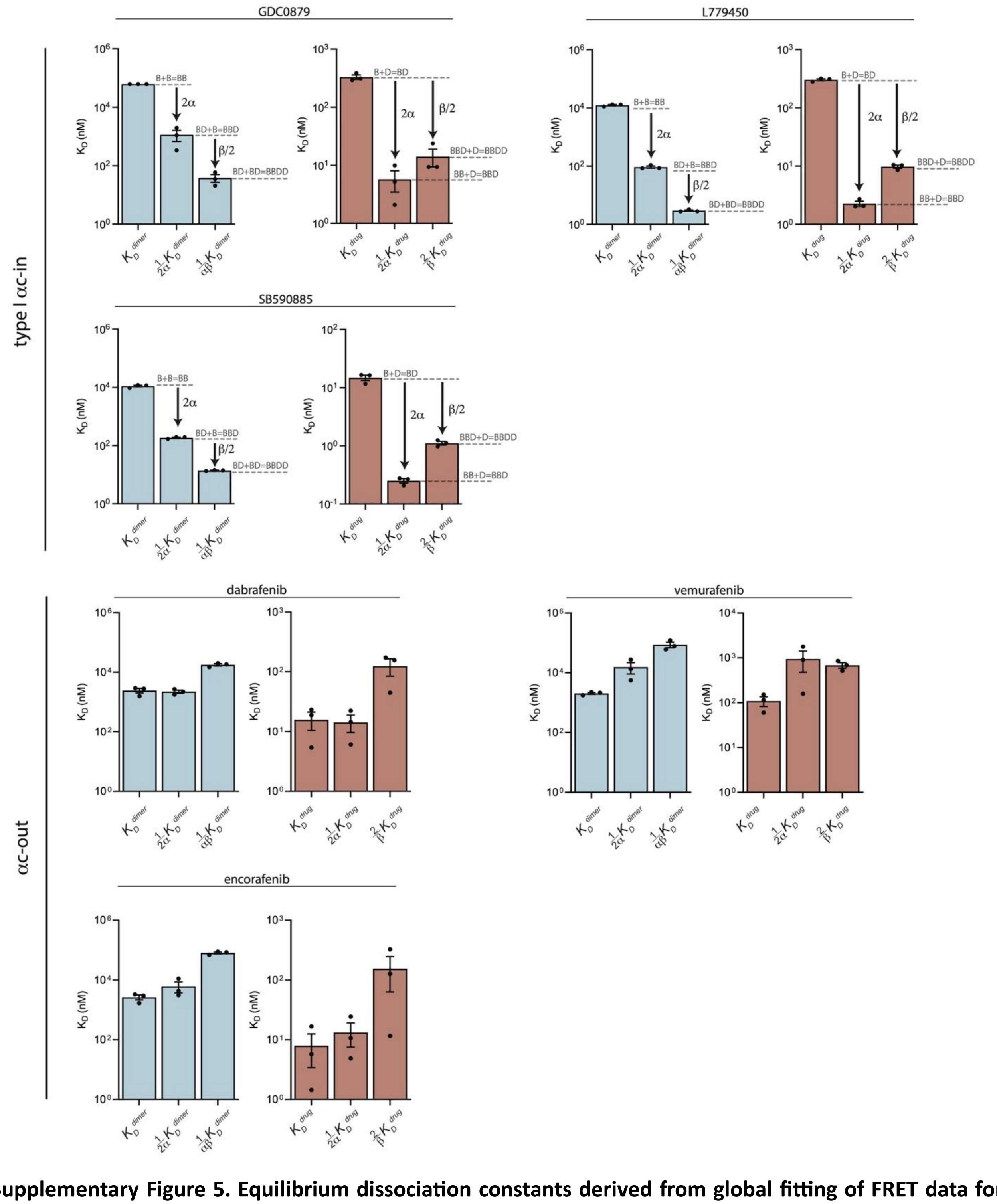


**Supplementary Figure 5. Equilibrium dissociation constants derived from global fitting of FRET data for all inhibitors.** Dissociation constants for BRAF dimerization (blue) and inhibitor binding affinity (red) derived from the global fitting analysis of the intermolecular FRET dimerization data. Allosteric coupling factors $\alpha$ and $\beta$ describe the coupling of the first and second inhibitor binding events to BRAF dimerization, respectively (see Methods). Similar values of $\alpha$ and $\beta$ for type I inhibitors indicate symmetric allosteric coupling, whereas for type II inhibitors the greater value of $\alpha$ compared to $\beta$ indicates asymmetric allosteric coupling. $\alpha$C-in type I and $\alpha$C-in type II both show a positive allosteric coupling and increase BRAF dimerization whereas $\alpha$C-out inhibitors show negative allosteric coupling and weaken $BRAF^{E586K}$ dimerization. Data represent best-fit values from global analysis of n=3 independent experiments performed in duplicate.

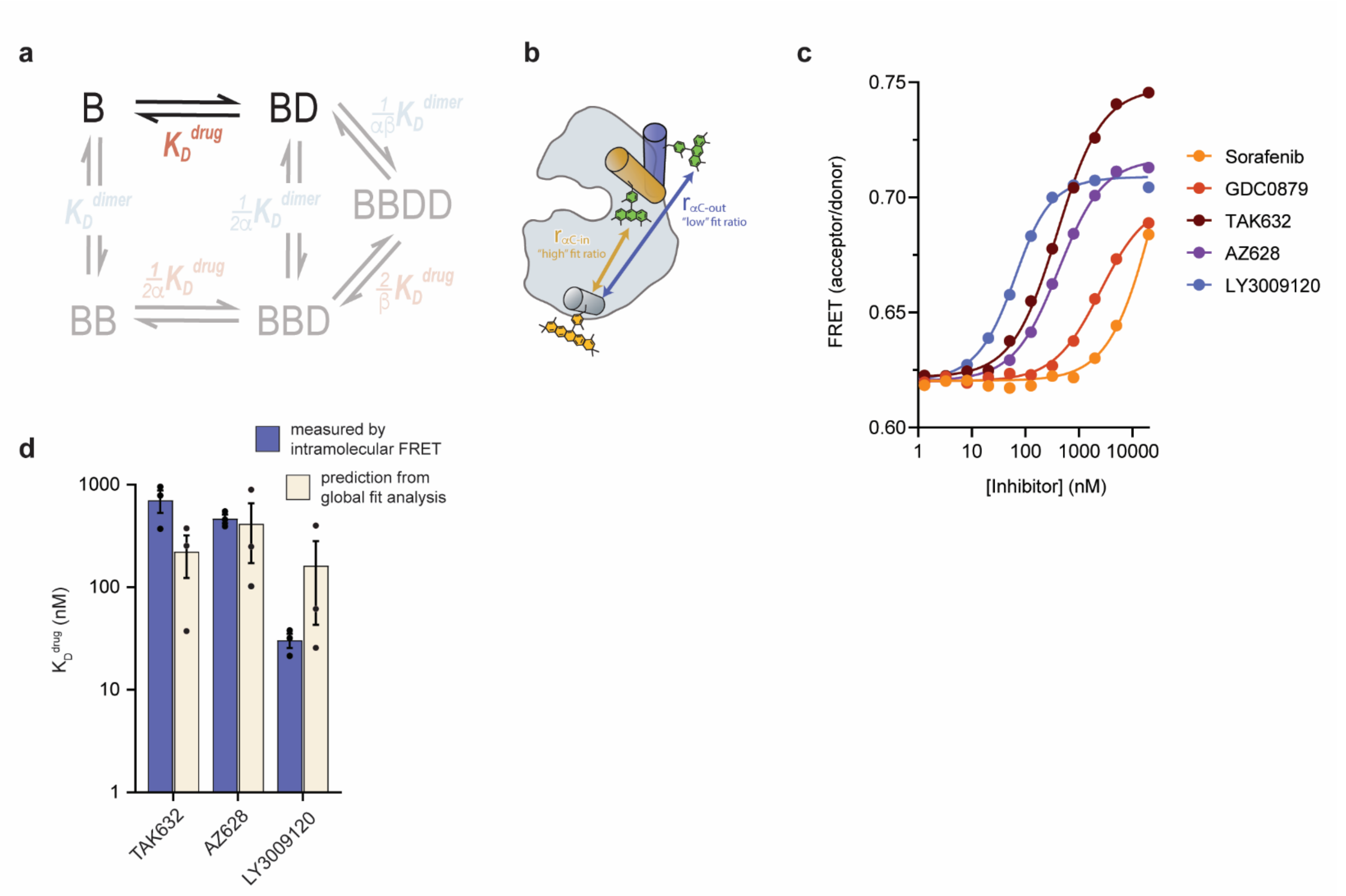


**Supplementary Figure 6. Inhibitor affinities for monomeric BRAF ($K_D^{drug}$) measured by intramolecular FRET. a)** Schematic highlighting the equilibrium dissociation constant that is being independently measured in the context of the thermodynamic model used for global fitting. **b)** Schematic showing the intramolecular FRET biosensor used to quantify inhibitor binding to monomeric BRAF containing dimer disrupting mutations. Inhibitor-induced conformational changes of the $\alpha$C-helix upon binding lead to a change in FRET. **c)** Inhibitor binding to monomeric BRAF as indicated by changes in FRET (acceptor / donor ratio). Binding curves were fit to a quadratic binding model in GraphPad Prism to extract $K_D^{drug}$. **d)** Best-fit values for $K_D^{drug}$ measured directly by intramolecular FRET (blue) in panel c compared to best-fit values predicted from the global fitting analysis of intermolecular FRET dimerization data (cream). Data represent the mean ± s.e.m.; n=3 independent experiments.

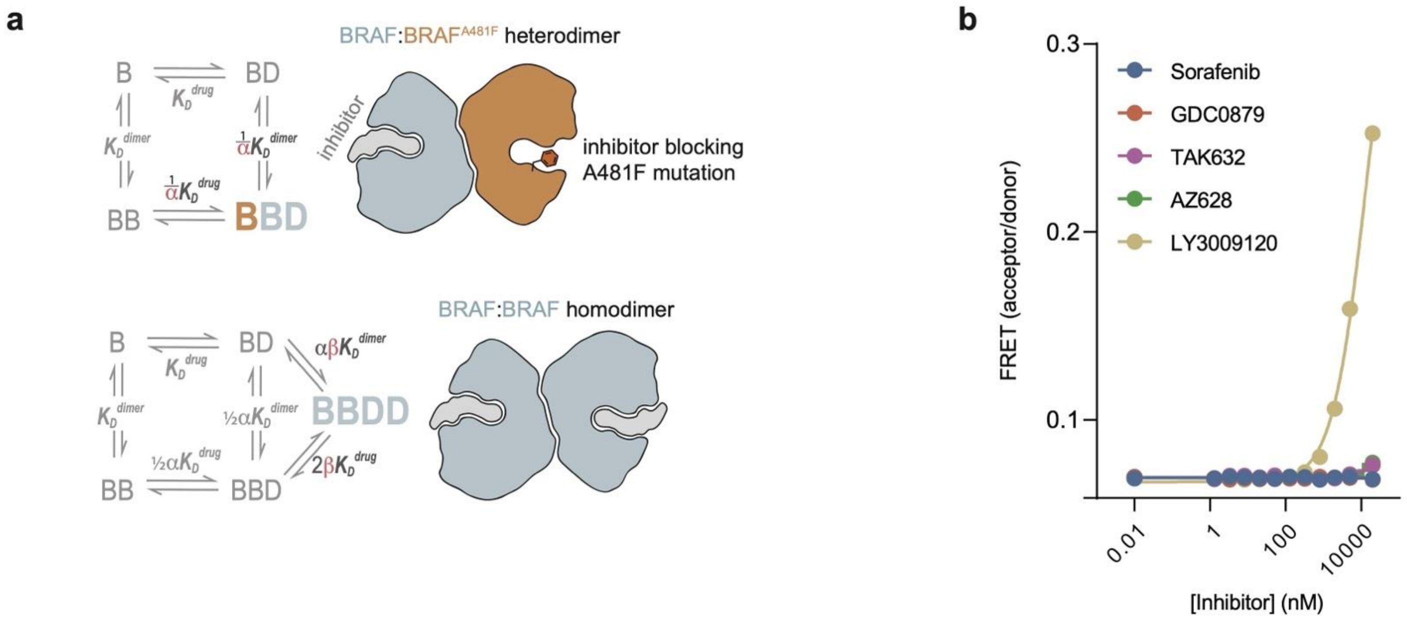


**Supplementary Figure 7. The A481F active site mutation blocks inhibitor binding and prevents inhibitor-induced dimerization. a)** A schematic demonstrating how the A481F active site mutation eliminates the effects of $\beta$ on BRAF dimerization by preventing the formation of BRAF dimers with both active sites bound to inhibitors. **b)** Intermolecular FRET dimerization data showing the formation of $BRAF^{A481F}$ / $BRAF^{A481F}$ homodimers as a function of $\alpha$C-in inhibitor concentration. $\alpha$C-in inhibitors fail to induce A481F homodimers indicating that the mutation effectively blocks inhibitor binding, with the exception of LY3009120 which, although it binds far more weakly to the mutant, does trigger dimerization at high concentrations.

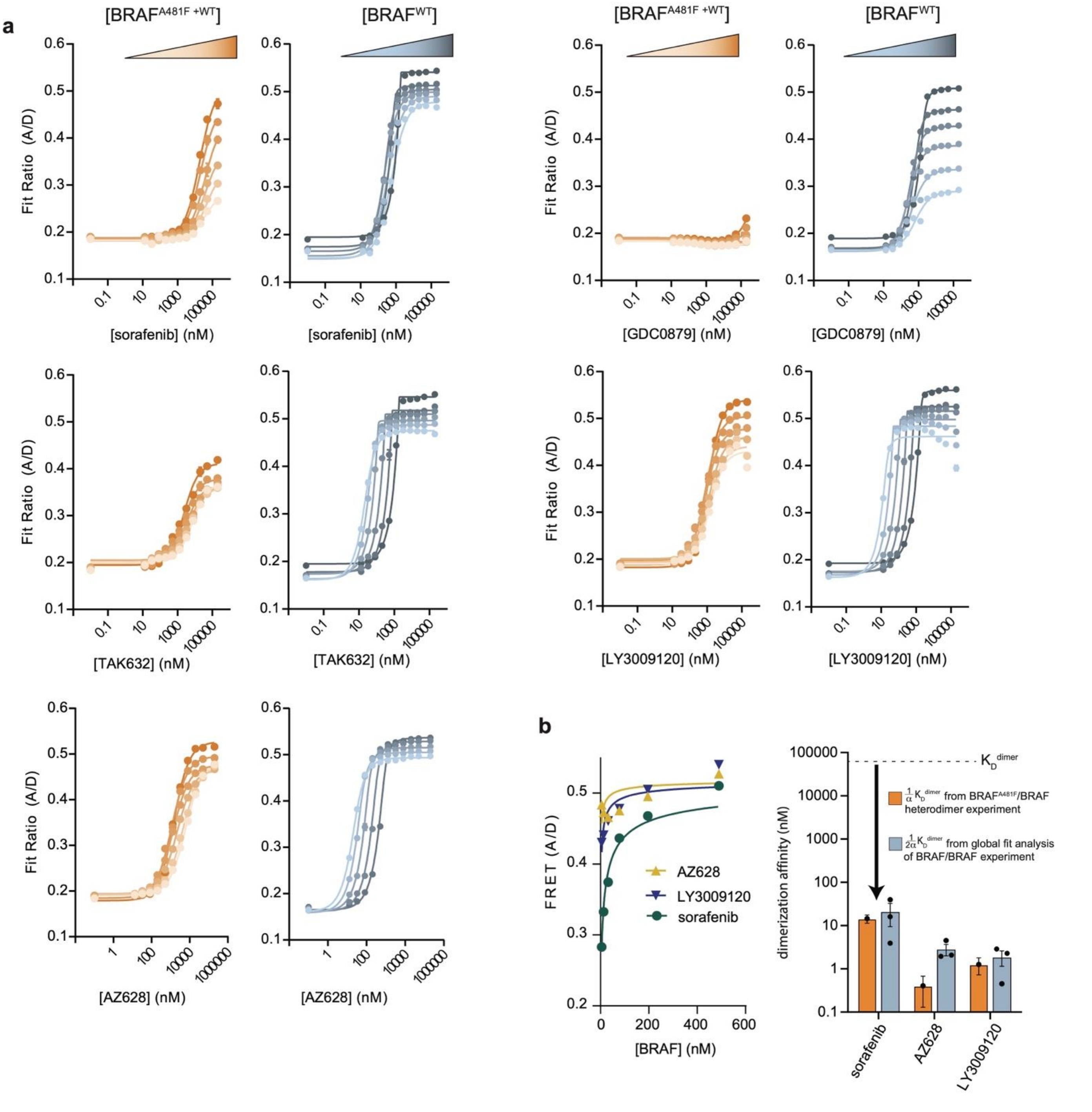


**Supplementary Figure 8. The binding of a single $\alpha$C-in type II molecule to the BRAF dimer is sufficient to dramatically increase dimerization affinity. a)** Representative intermolecular FRET dimerization data showing the formation of BRAF$^{A481F}$/BRAF heterodimers (orange) and BRAF/BRAF homodimers (blue) as a function of inhibitor concentration. All type II inhibitors strongly induce BRAF$^{A481F}$/BRAF heterodimers, while the type I inhibitor GDC0879 does not. **b)** The upper baselines observed at saturating inhibitor concentrations in panel a were fit to a monomer-dimer equilibrium model (left) to obtain estimates of the $K_D$ values for saturated BBDD dimers (right), which agree well with $\frac{1}{2\alpha}K_D^{dimer}$ values calculated from the global fitting analysis of independent FRET experiments.

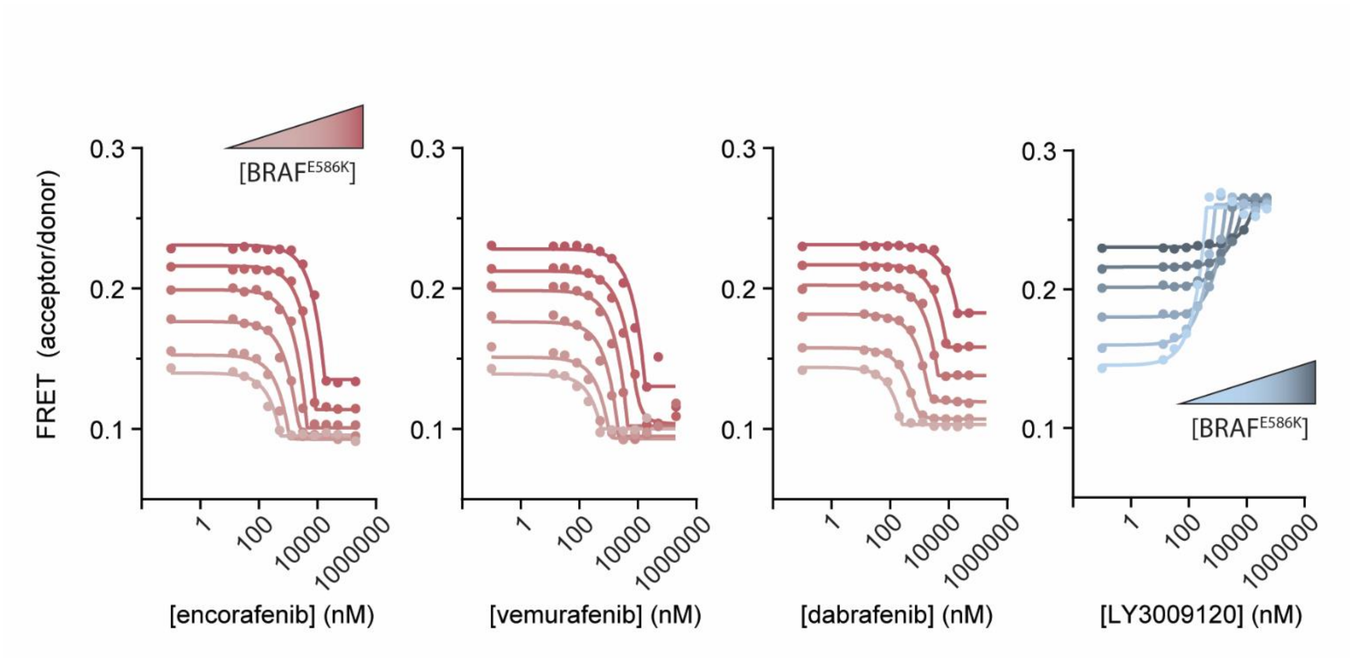


**Supplementary Figure 9. Intermolecular FRET experiments measuring the disruption of BRAF$^{E586K}$ dimerization by $\alpha$C-out inhibitors.** Representative data showing BRAF$^{E586K}$ dimerization as a function of $\alpha$C-out inhibitors (red) and the $\alpha$C-in inhibitor LY3009120 for comparison. Gradients from light to dark represent experiments done at increasing BRAF concentration. Lines are fits of individual titrations to a quadratic binding model in Graphpad prism and are for illustrative purposes only; all data were globally fit to the thermodynamic model shown in Figure 1c.

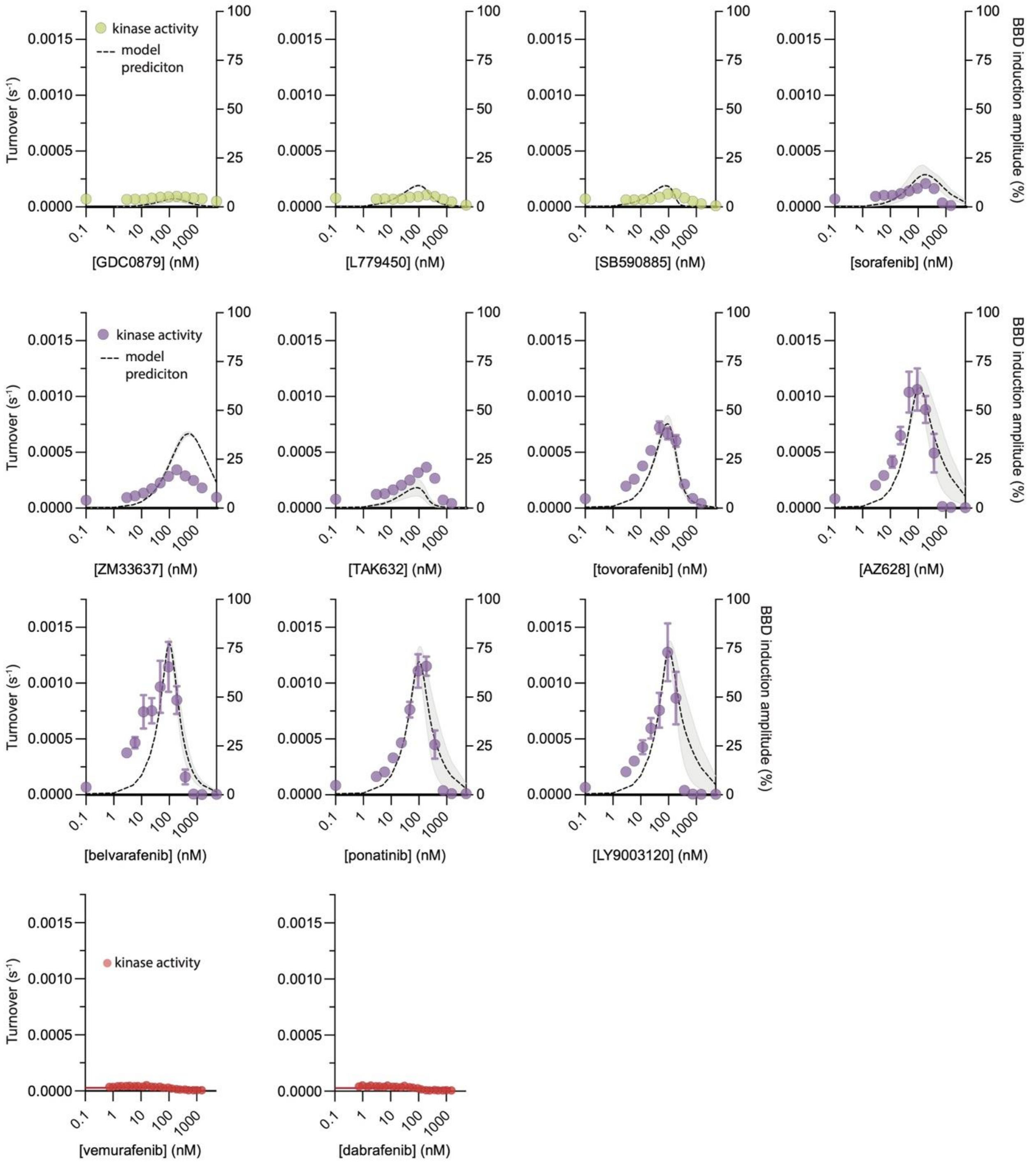


**Supplementary Figure 10. Simulations of partially occupied BRAF dimer formation and *in vitro* kinase activity.** The formation of partially occupied "BBD" BRAF dimers was simulated as a function of inhibitor concentration using the allosteric model (Figure 1c) parameterized for each inhibitor via the global analysis of intermolecular FRET data (see Methods). Multiple independent simulations were performed for each inhibitor based on models parameterized with separate FRET datasets and reported as BBD induction percent. The mean of these simulations from n=3 independent experiments is shown as a black dotted line with the ± s.e.m. shown as a light gray band. In vitro phosphorylation of MEK by BRAF was measured as a function of $\alpha$C-in type I (yellow), $\alpha$C-in type II (purple), and $\alpha$C-out (red) inhibitors and overlayed onto the BBD induction amplitude simulations. The inhibitor-induced BRAF activation by the asymmetrically coupled type II inhibitors is far greater than the symmetrically coupled type I inhibitors, in excellent agreement with the allosteric model simulations. Data represent the mean ± s.e.m.; n=3 independent experiments performed in duplicate.

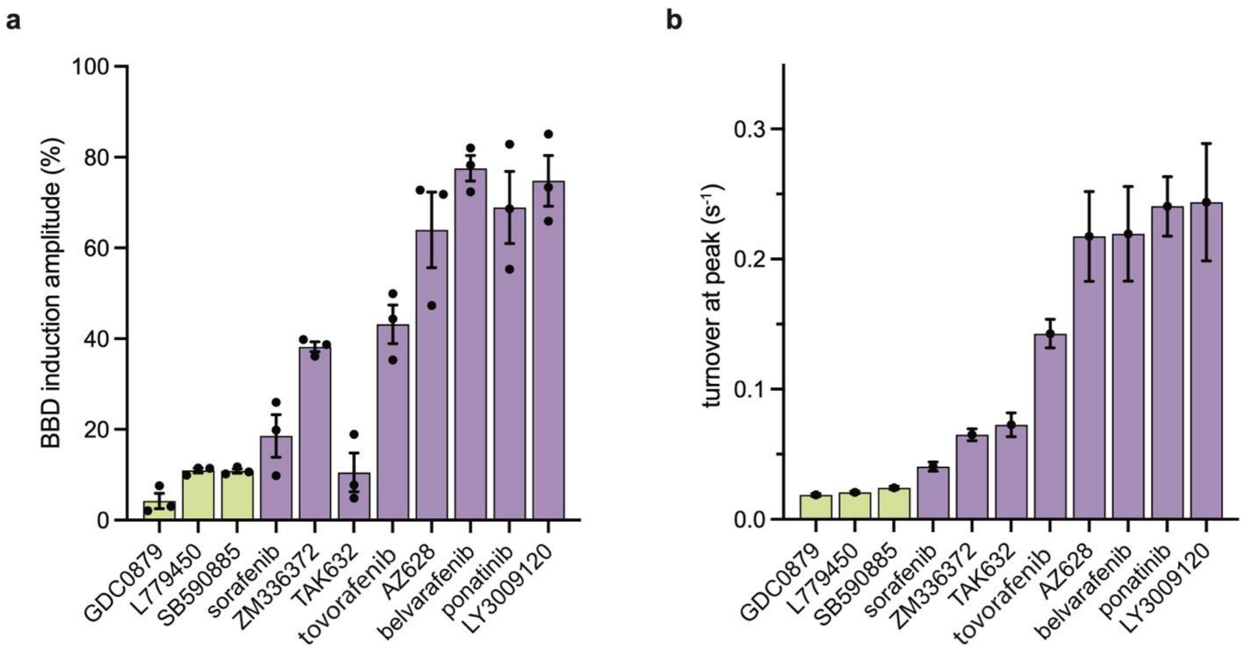


**Supplementary Figure 11. BBD induction amplitude and peak catalytic turnover. a)** Peak BBD induction amplitudes from individual simulations shown in Supplementary Fig. 10 are shown for type I (yellow) and type II (purple) $\alpha$C-in inhibitors. Data represent the mean ± s.e.m.; n=3 independent experiments performed in duplicate. **b)** Peak inhibitor-induced BRAF kinase activity for $\alpha$C-in inhibitors determined by in vitro kinase activity assays measuring MEK phosphorylation by BRAF shown in Supplementary Fig. 10. Data represent the mean and 95% CIs of peak kinase activity determined by fitting n=3 independent kinase activity experiments done in duplicate to a bell-shaped dose-response model in GraphPad Prism.

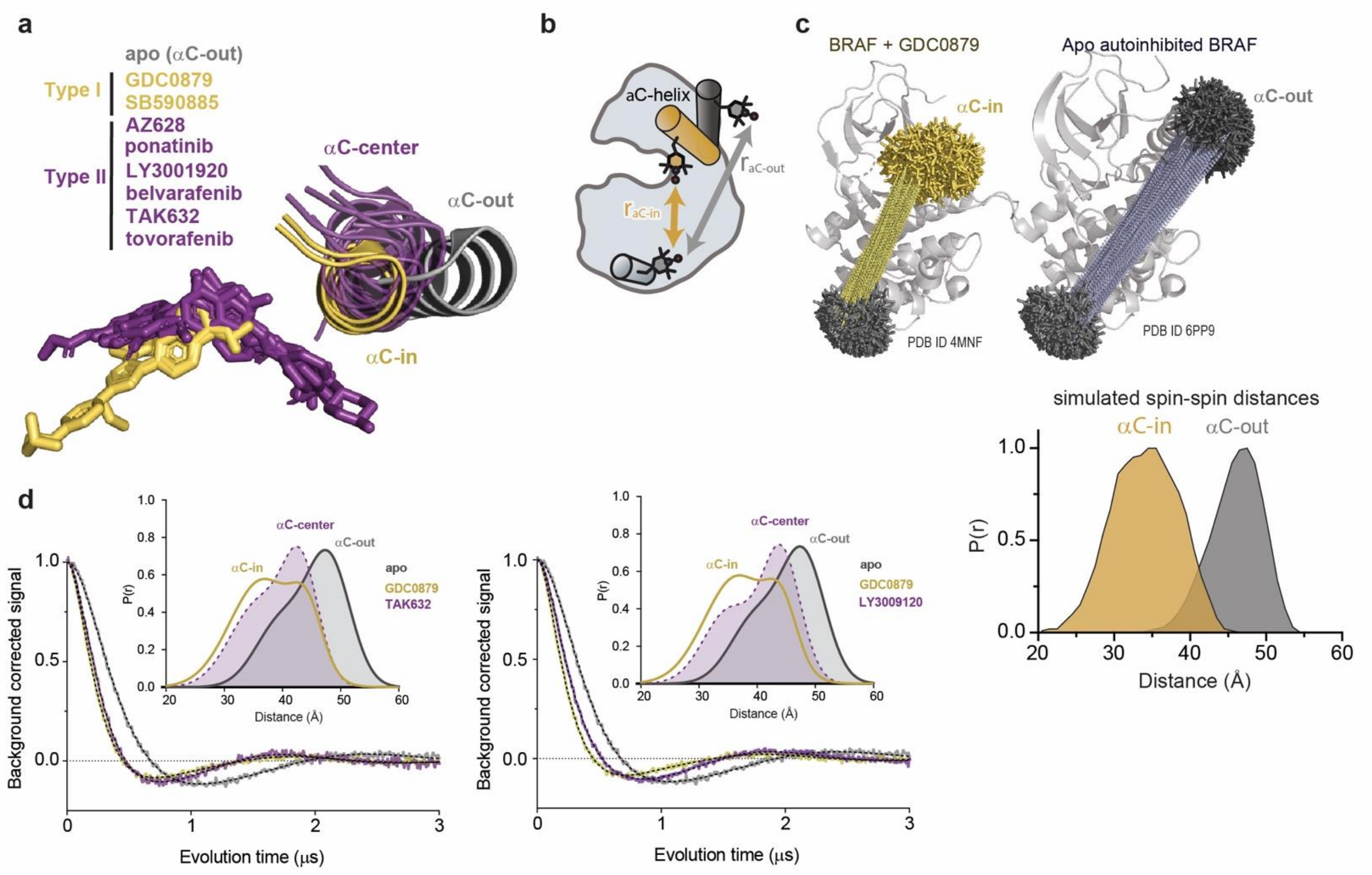


**Supplementary Figure 12. Measuring the conformation of the αC-helix with DEER. a)** X-ray structures of BRAF in the apo state (grey) (PDB ID: 6PP9), bound to type I inhibitors (yellow) (PDB IDs: 4MNF, 2FB8) and type II inhibitors (purple) (PDB IDs: 4RZW, 6P3D, 5C9C, 6XFP, 4KSP, 6V34) highlighting the positioning of the αC-helix. Structures were aligned on the C-terminal lobe of the kinase. **b)** Schematic of the labeling strategy used to track the conformation of the αC-helix. **c**) MtsslWizard was used to generate spin-spin distance distributions using BRAF crystal structures. Rotamers of 4-maleimido-TEMPO were generated from a GDC0879-bound BRAF structure (PDB ID: 4MNE) adopting an αC-in state (yellow) and a structure of apo BRAF (PDB ID: 6PP9) adopting an αC-out state (grey). Individual spin-spin distances (dashed lines) were calculated and compiled to simulate spin-spin distances used to assign the Gaussian distance distributions from DEER experiments measuring the αC-helix represented in panel d. **d)** DEER waveforms and Gaussian distance distributions for apo BRAF and BRAF bound to type I inhibitor GDC0879 (yellow) and type II inhibitors TAK632 and LY3009120 (purple).

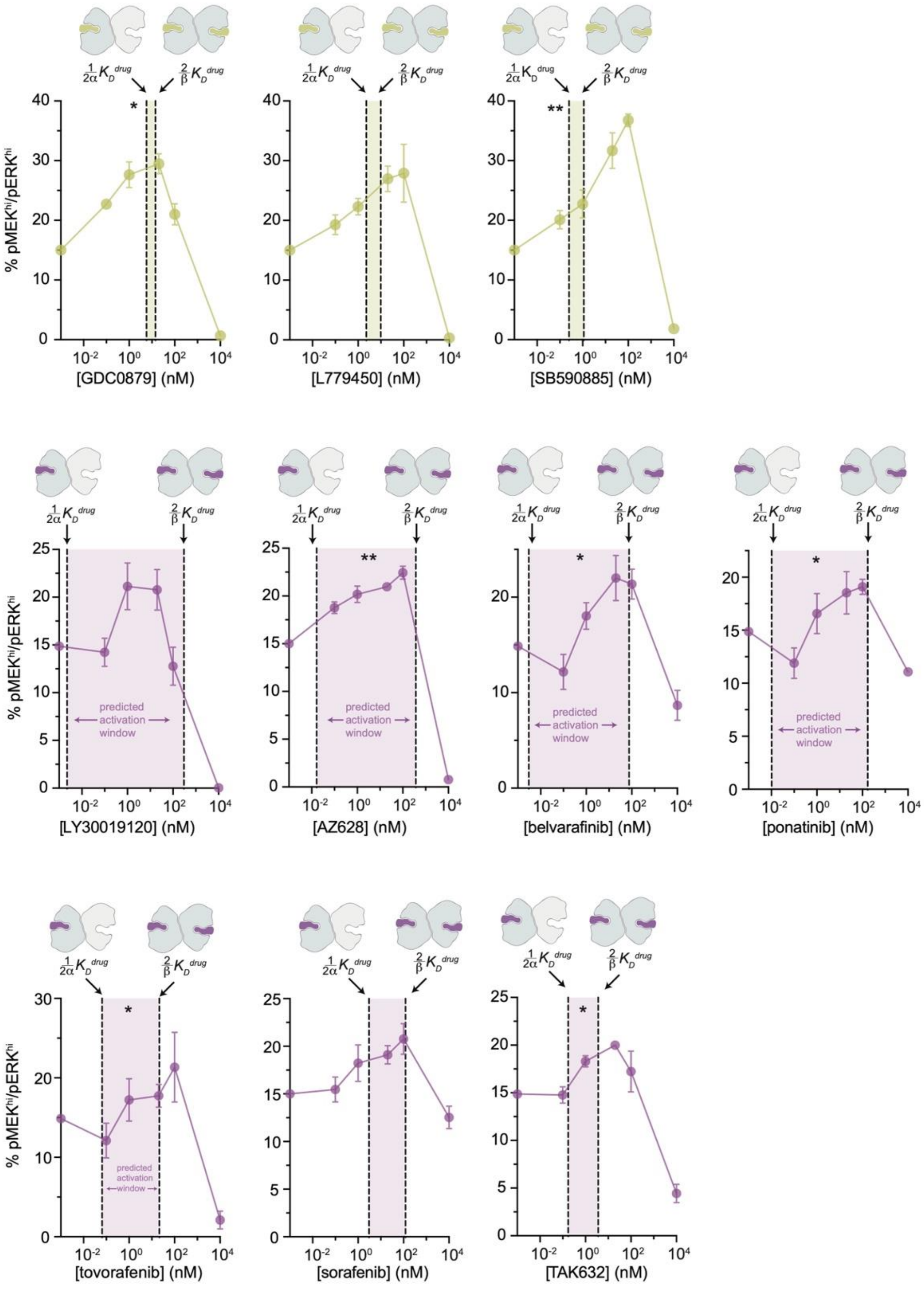


**Supplementary Figure 13. Inhibitor-induced MAPK/ERK pathway activation in SK-MEL-2 cells.** Paradoxical activation of MAPK/ERK signaling in SK-MEL-2 cells by type I (yellow) and type II (purple) RAF inhibitors. The dashed lines represent the inhibitor affinities for the first and second subunit of the BRAF dimer ($\frac{1}{2\alpha}K^{drug}$ and $\frac{2}{\beta}K^{drug}$, respectively), determined from the global fitting analysis with the shaded gap between the two lines representing the predicted window of activation. Error bars: mean ± s.e.m.; n=3. Significance: 1-way ANOVA/Geisser-Greenhouse/Tukey's test. Asterisks reflect significant paradoxical activation between pairs of data points for GDC0879 [0→0.1 nM,*p=0.0105], [0→20*p=0.0459]; SB590885 [0→100**p=0.0085], [0.1→100**p=0.0064]; AZ628 [0→20**p=0.0036], [0→100*p=0.0256]; belvarafenib [0.1→1*p=0.0250], [0.1→100*p=0.0105]; ponatinib [1→20*p=0.0135]; tovorafenib [0.1→1*p=0.0328]; and TAK632 [0→20*p=0.0249], [0.1→20*p=0.0361]. Most pairwise comparisons [x→10000 nM] also rose to significance and comparisons not mentioned did not.

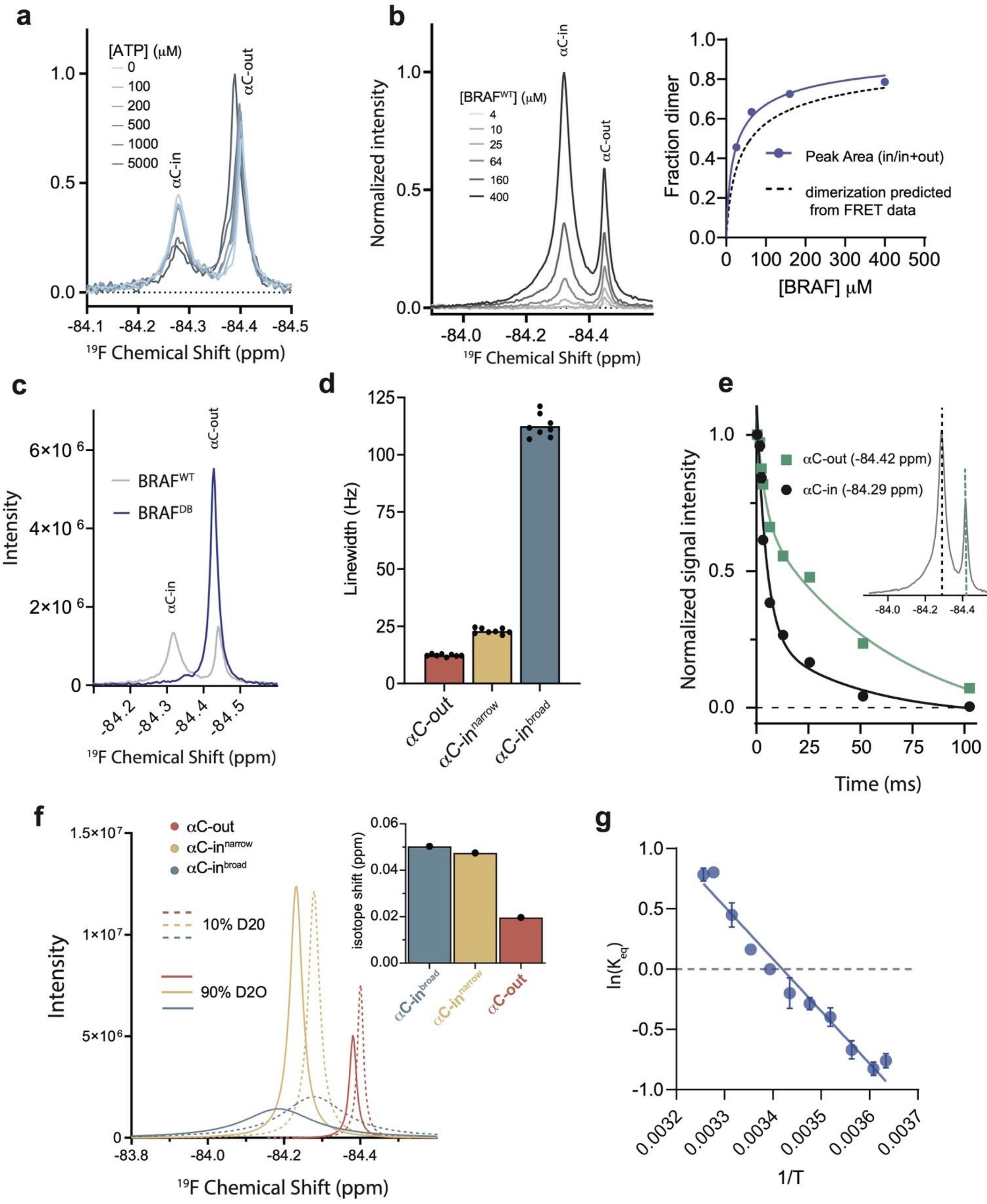

a
[ATP] (μM)
0
100
200
500
1000
5000
αC-in
αC-out
¹⁹F Chemical Shift (ppm)
b
Normalized intensity
[BRAF^WT] (μM)
4
10
25
64
160
400
αC-in
αC-out
¹⁹F Chemical Shift (ppm)
Fraction dimer
Peak Area (in/in+out)
dimerization predicted from FRET data
[BRAF] μM
c
Intensity
BRAF^WT
BRAF^DB
αC-in
αC-out
¹⁹F Chemical Shift (ppm)
d
Linewidth (Hz)
αC-out
αC-in^narrow
αC-in^broad
e
Normalized signal intensity
αC-out (-84.42 ppm)
αC-in (-84.29 ppm)
Time (ms)
f
αC-out
αC-in^narrow
αC-in^broad
10% D20
90% D2O
Intensity
isotope shift (ppm)
αC-in^broad
αC-in^narrow
αC-out
¹⁹F Chemical Shift (ppm)
g
ln(K_eq)
1/T

**Supplementary Figure 14. $^{19}$F NMR resonance assignments and experiments confirming the presence of dynamic heterogeneity within the BRAF dimer. a)** $^{19}$F NMR spectra of BRAF labeled on the $\alpha$C-helix with BTFA in the presence of AZ628 (purple) and ATP (blue). AZ628 causes a decrease in the upfield resonance and increase in the downfield resonance, whereas ATP causes the opposite effect, allowing the upfield resonance to be assigned to the monomer/$\alpha$C-out state and the downfield resonance to be assigned to the dimer/$\alpha$C-in state. **b)** $^{19}$F NMR spectra of BRAF at different BRAF concentrations. The relative peak areas of the $\alpha$C-in peak versus the $\alpha$C-out peak were plotted as a function of BRAF concentration (upper right) and fit to a monomer-dimer equilibrium model (blue), to derive a value for the dimerization affinity which was in reasonable agreement with the value determined from the FRET experiments (dotted line) with a $K_D^{dimer}$ of 32.2 $\pm$ 9.4 $\mu$M. **c)** $^{19}$F NMR spectra of BRAF (gray) and BRAF containing dimer disrupting mutations (blue). The disappearance of the downfield resonance in the presence of the dimer disrupting mutations is consistent with the resonance assignments discussed in **a** and supports the assignment of the exchange broadened peak to the dimer state. **d)** Linewidths extracted from deconvoluted $^{19}$F NMR spectra corresponding to the individual species represented in Figure 4a. Data represent the best-fit values from the spectral deconvolution of n=8 independent experiments. **e)** $^{19}$F NMR $T_2$ relaxation profiles obtained from BRAF. Intensities at -84.42 ppm (green) and -84.29 ppm (black) corresponding to the $\alpha$C-out and $\alpha$C-in resonances, respectively, were plotted as a function of decay time. Relaxation profiles required a multi-exponential fit ($p<0.0001$) indicating the presence of overlapping resonances with distinct relaxation times. Data represent the mean ± s.e.m.; n=15 independent experiments. **f)** Solvent isotope effect experiments obtained with BRAF in 10% $D_2O$ (dotted lines) and 90% $D_2O$ (solid lines). Individual component fits from spectral deconvolution are shown and correspond to the $\alpha$C-out (red), $\alpha$C-in$^{narrow}$ (yellow), and $\alpha$C-in$^{broad}$ (blue) states. Relative peak shifts indicate that the two $\alpha$C-in states exhibit a similar degree of solvent exposure. **g)** Van ’t Hoff plot showing the temperature dependence of the equilibrium between the narrow and broad dimer states.

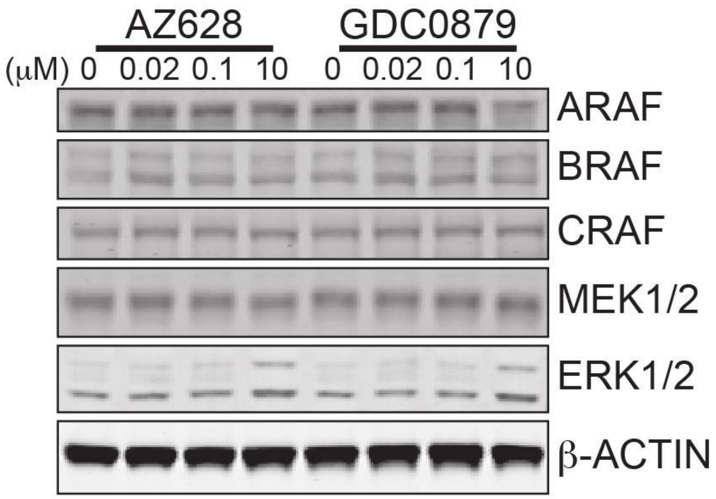


**Supplementary Figure 15. Western blot confirming the presence of all RAF isoforms in SK-MEL-2 cells.** SK-MEL-2 cells were treated with the indicated amount of type II inhibitor AZ628 or type I inhibitor GDC0879 for 1 hr. Cell lysates were immunoblotted for all endogenous RAF isoforms. Note that ARAF, BRAF, and CRAF are present at similar levels.

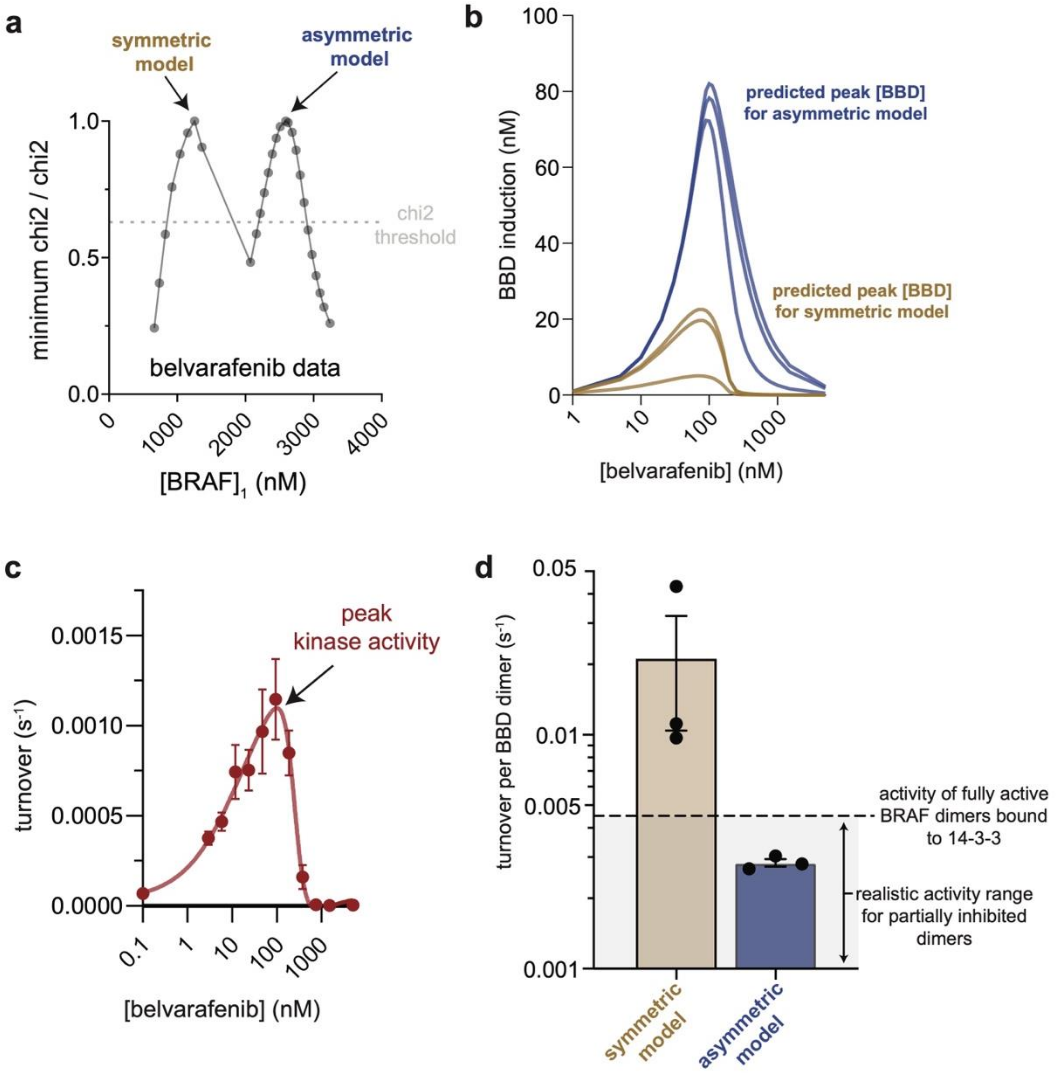


**Supplementary Figure 16. Only asymmetric allosteric models correspond to realistic levels of catalytic activity of BBD dimers. a)** For some datasets, removing constraints on the BRAF concentrations used in the global fitting analysis resulted in two possible solutions, as shown for a representative case of fitting FRET data obtained with belvarafenib, where a one-dimensional error surface for the [BRAF]$_1$ parameter reveals two minima. The grey dotted line represents the chi$^2$ threshold used to establish the 95% CI for the [BRAF]$_1$ parameter. The two minima at different BRAF concentrations with equivalent chi$^2$ values correspond to a symmetric and an asymmetric model. **b)** The induction of partially occupied "*BBD*" BRAF dimers was simulated as a function of belvarafenib concentration using either the symmetric (brown) or asymmetric (blue) allosteric models. Simulations were performed for three independent FRET data sets. **c)** Peak kinase activity for belvarafenib was determined by fitting BRAF kinase activity to a bell-shaped dose-response curve. **c)** Peak kinase activity and predicted peak [BBD] were used to calculate the inferred catalytic turnover per BBD dimer from the two models. Values were compared to the previously reported catalytic activity of 14-3-3-bound BRAF dimers (black dotted line)[1]. Note that the inferred activity of BBD dimers predicted by the symmetric model (brown) is far higher than the published value for fully activated and uninhibited BRAF dimers. Conversely, the catalytic activity predicted by the asymmetric model (blue) is in close agreement with the published value and is slightly lower, consistent with one of the two active sites of the dimer being inhibited.

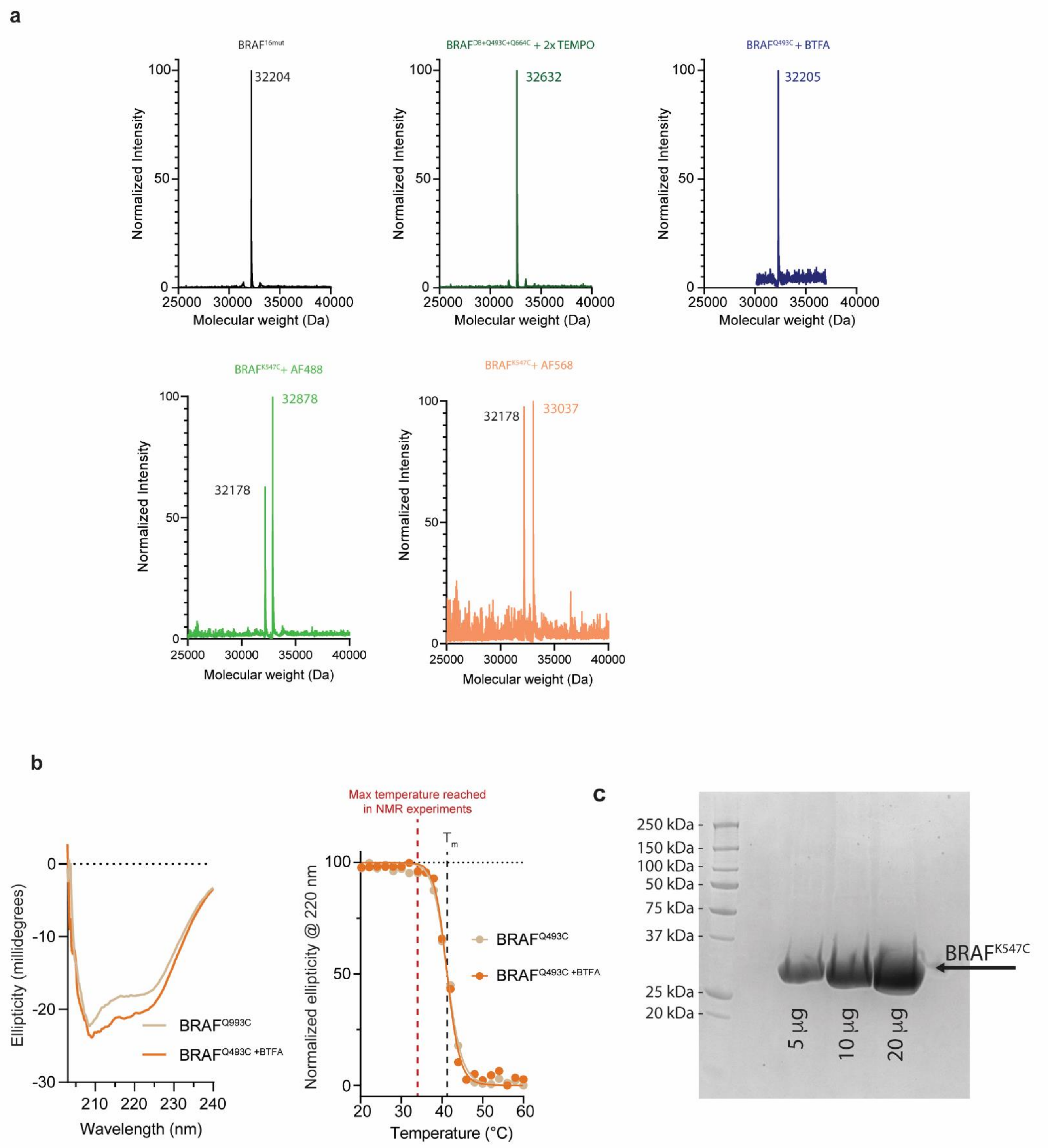


**Supplementary Figure 17. Validation of BRAF labeling, stability, and purity. a)** Mass spectra of BRAF$^{16mut}$ (theoretical molecular weight: 32,204 Da), BRAF$^{DB+Q493C+Q664C}$ labeled with two 4-Maleimido-TEMPO spin probes (theoretical molecular weight: 32,638 Da), BRAF$^{Q493C}$ labeled with 3-Bromo-1,1,1-trifluoroacetone (BTFA, theoretical molecular weight: 32,205 Da), BRAF$^{K547C}$ (32,178 Da) labeled with Alexa Flour 488 (theoretical molecular weight: 32,900 Da), BRAF$^{K547C}$ (32,178 Da) labeled with Alexa Fluor 568 ( theoretical molecular weight: 33,060 Da. **b)** Circular dichroism spectra of BRAF$^{Q493C}$ (tan) and BRAF$^{Q493C}$ labeled with BTFA (orange). Ellipticity at 220 nm was plotted as a function of temperature. Note that the presence of the BTFA probe does not influence the melting temperature ($T_m$) which is well above the maximum temperature used in NMR experiments. **c)** SDS-PAGE gel of BRAF$^{K547C}$ at increasing total protein amounts showing the purity of the samples used in intermolecular FRET experiments.

| inhibitor | source | monoisotopic mass [M+Na]+ (Da) | measured mass [M+Na]+ (Da) | |
|---|---|---|---|---|
| vemurafenib | Selleck Chemicals | 512.06178 | 512.0394 | |
| encorafenib | Selleck Chemicals | 562.14101 | 562.1418 | $\alpha$C-out |
| Dabrafenib | Selleck Chemicals | 542.0903 | 542.0917 | |
| GDC0879 | Selleck Chemicals | 357.1322 | 357.1321 | |
| SB590885 | TargetMol | 476.20571 | 476.204 | $\alpha$C-in type I |
| L779450 | Selleck Chemicals | 348.0898 | 348.0895 | |
| Sorafenib Tosylate | Selleck Chemcials | 465.0936 | 465.093 | |
| TAK632 | Selleck Chemcials | 577.0928 | 577.0924 | |
| AZ628 | Selleck Chemcials | 474.19 | 474.1904 | |
| LY3009120 | Selleck Chemcials | 447.2279 | 447.2273 | $\alpha$C-in type II |
| Ponatinib | TargetMol | 555.2091 | 555.2087 | |
| ZM336372 | Selleck Chemcials | 412.1632 | 412.1631 | |
| Belvarafenib | Selleck Chemcials | 501.0671 | 501.0675 | |
| Tovorafenib | Selleck Chemcials | 527.99947 | 528.0014 | |

**Supplementary Table I. Vendor sources and validation by mass spectrometry.**
